# Strong signatures of selection on genes underlying core reinforcement mechanisms in speciating desert tortoises

**DOI:** 10.1101/2024.06.06.597788

**Authors:** Sarah M. Baty, Raúl Araya-Donoso, Avery Paulsen, Avery Williams, Dale F. DeNardo, Kenro Kusumi, Greer A. Dolby

## Abstract

Genomic reinforcement and differential ecological adaptation are thought to be fundamental mechanisms of speciation. In this study we investigate the genomic basis of adaptation and reinforcement between two desert tortoise species of North America that occupy desert habitats with differing seasonal rainfall patterns and have considerable behavioral and reproductive differences yet maintain a narrow hybrid zone. We generated a chromosome-scale reference genome for *Gopherus morafkai* and performed analysis of synteny, genes under positive selection, and environmental niche modeling. Results show extensive positive selection (422 genes) including related to eye development and function that may relate to environmental differences, as well as prezygotic isolation mechanisms such as sperm-egg recognition, and postzygotic reinforcement mechanisms such as the spindle assembly checkpoint, and sister chromatid pairing. Together, results offer strong genetic support for the role of these classic processes in shaping reproductive isolation and lineage divergence of speciating tortoises.

## INTRODUCTION

Environmental and ecological disparities are known to facilitate ecological speciation^1^; examples of this include contrasting magnitudes and timing of precipitation or unique vegetation assemblages^2,3^. While ecological adaptations may differentiate species, the maintenance of lineage divergence and speciation requires the accumulation of reinforcement mechanisms in the form of prezygotic and postzygotic barriers that aid reproductive isolation. Examples of genomic reinforcement mechanisms include meiotic and mitotic incompatibilities^4^, chromosomal changes that prevent proper pairing and segregation of chromosomes^5^, and reproductive changes affecting sperm-egg recognition and fertilization^6^. Whole genome analysis offers an opportunity to understand how these molecular mechanisms arise in nature and empirically test predictions from reinforcement theory.

Desert tortoises *Gopherus morafkai* and *G. agassizii* diverged ∼4–5 million years ago^7–9^, reside in environments with differing seasonal rainfall patterns, and are thought to have undergone fast, parapatric or allopatric speciation due at least in part to this ecological difference^10,11^. The Mojave Desert receives less annual rainfall, which typically occurs only in winter months^12,13^. In contrast, the Sonoran Desert receives considerable rainfall during the summer monsoon season and intermediate levels of winter rainfall^13–15^. It is thought that the Colorado River reduces gene flow between the species but does not entirely prevent it^16^, as evidenced by a narrow hybrid zone east of the Colorado River in the ecotone between the Mojave and Sonoran deserts. The seasonal differences in precipitation also drive species’ differences in foraging behaviors and activity patterns^16,17^. In addition to the precipitation gradient, the Mojave to Sonoran habitat transition presents a gradient in UV index^18^, potentially serving as a driver of differences in winter basking behavior between the species^19^.

*Gopherus morafkai* and *G. agassizii* also exhibit differences in morphology^8,11^, clutch size and frequency^20^, brumation behavior^19–21^, body size^22^, and lifespan^22^. Yet, the hybrid zone between these species indicates they have not yet reached complete reproductive isolation, offering an opportunity to assess to what degree genomic reinforcement mechanisms have developed in this lineage pair. To do this we developed a chromosome-scale reference genome for *G. morafkai* and performed comparative genomic analyses with the published reference genome for *G. agassizii*^23^. We analyzed synteny, genes under selection and their biological enrichment and interactions, as well as habitat modeling to assess the underlying genomic mechanisms underlying ecological differences and genomic reinforcement mechanisms between these desert tortoises.

## RESULTS

### Genome Assembly and Annotation

The *G. morafkai* genome was assembled using PacBio and Illumina sequencing and Hi-C scaffolding technologies. The assembly size is 2.2 Gb with 1,557 scaffolds. The N50 of the assembly is 14.7 Mb and the L50 is 47. The genome assembly had 248 (97.3%) complete BUSCOs, with 2 fragmented BUSCOs, (0.8%) and 5 missing orthologs. The predicted transcripts generated from the genome annotation had 200 complete BUSCOs (78.5%) with only 39 fragmented BUSCOs (15.3%) and 16 missing orthologs (6.2%). We annotated 22,130 protein coding gene models with an average gene length of 19,422 bp, which is comparable to the *G. agassizii* annotation that has 25,469 predicted protein coding genes.

### Analysis of Synteny Conservation

To identify changes in shared synteny between *Gopherus* and other turtles we analyzed gene order conservation between *G. morafkai* and *Trachemys scripta elegans*, the red-eared slider (Figure 1). When mapped to the *G. morafkai* reference, *T. s. elegans* shows several small gene order rearrangements on scaffolds representing chromosomes 2, 9 13, and 14. There are larger chromosomal rearrangements occurring on scaffold 16, mapping to segments from three different representative scaffolds (21, 22, and 23) from *G. morafkai*. These chromosomal rearrangements could have contributed to the divergence of *Gopherus* from the ancestral chelonians. Pairwise synteny analysis between *G. morafkai* and *G. agassizii* showed high synteny conservation, with a scaffold fusion in the *G. agassizii* genome on scaffold 24 (scaffolds 24 and 25 in *G. morafkai*; Supplementary Figure 4). We estimate this to be an assembly artefact of the *G. agassizii* reference based on a mis-join from Chicago scaffolding data as this join occurs in a poly-N region, identified through scaffold analysis with a resolution of around 1 kbp.

**Figure 1:**
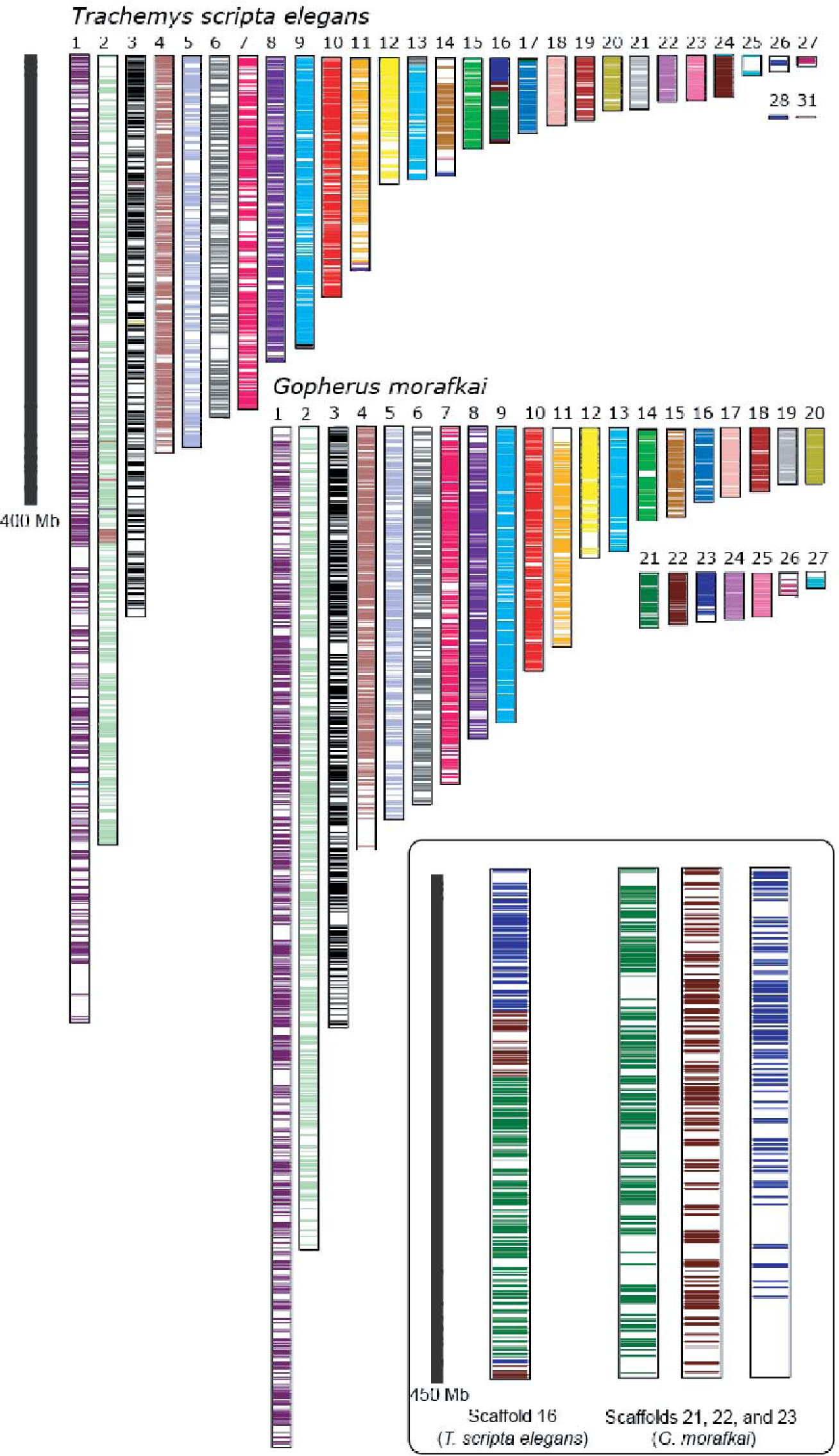
Synteny comparisons across the most gene rich scaffolds in the genome assemblies of *G. morafkai* and *T. s. elegans*. Scaffold 16 of the *T. s. elegans* assembly shows fusion of three scaffolds of the *G. morafkai* assembly from 21, 22 and 23. See Supplemental Figure 4 for synteny comparisons between *G. morafkai* and *G. agassizii*.

### Ecological Niche Analysis

A principal components analysis (PCA) using bioclimatic variables representative of yearly climatic variation was performed to characterize the climatic conditions of *G. agassizii* and *G. morafkai* habitats (Figure 2C). The first principal component PC1 positively correlated with annual precipitation and negatively with temperature seasonality, whereas PC2 negatively associated with mean annual temperature and other temperature variables. Significant niche divergence was detected between the tortoises with a niche overlap of 0.026 (p < 0.001). Although there is some level of niche overlap where the two species interact and hybridize, *G. agassizii* resides in the drier environments with highly seasonal temperatures, and *G. morafkai* occurs in wetter areas with less seasonal variation in temperatures.

**Figure 2:**
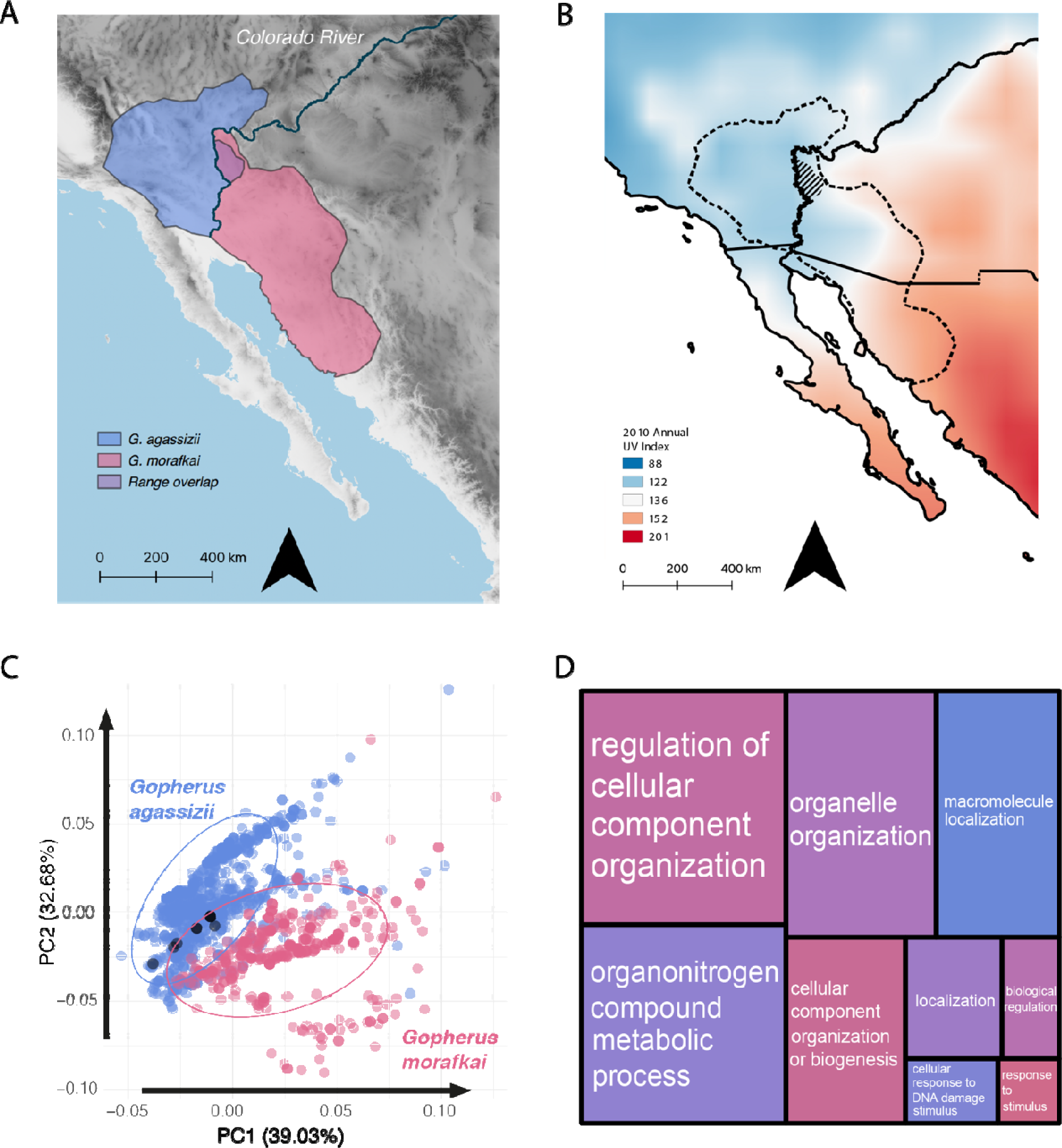
Summary of habitat and genomic differences. A) Range maps of *G. morafkai* and *G. agassizii;* B) UV index across the North American deserts; C) a PCA of environmental niche models for *G. morafkai* and *G. agassizii*. Black circles are locality points within the hybrid zone of the two species. D) REVIGO treemap of the significantly enriched biological processes based on genes under selection between *G. morafkai* and *G. agassizii.* The size of the treemap rectangles are representative of the p value associated with each term. See Supplemental Figure 6 for the detailed categories within each rectangle from D.

### Functional Enrichment and Interaction of Genes under Positive Selection

To characterize potential differential adaptations in protein coding genes of the two species, we calculated pairwise dN/dS. This yielded 422 genes under significant positive selection (hereafter, PS), 389 of which were annotated in the *G. morafkai* annotation. PS genes were enriched in biological processes including *biogenesis*, *localization*, *biological regulation, cellular response to DNA damage stimulus, and response to stimulus* (Figure 2D; see Table 3 for full list). The interactions among positively selected genes according to STRING showed significant enrichment values, indicating that the genes under PS interact with one another more than expected by chance (Figure 3). The enrichment p value was 0.0267 and average local clustering coefficient of 0.33.

**Figure 3:**
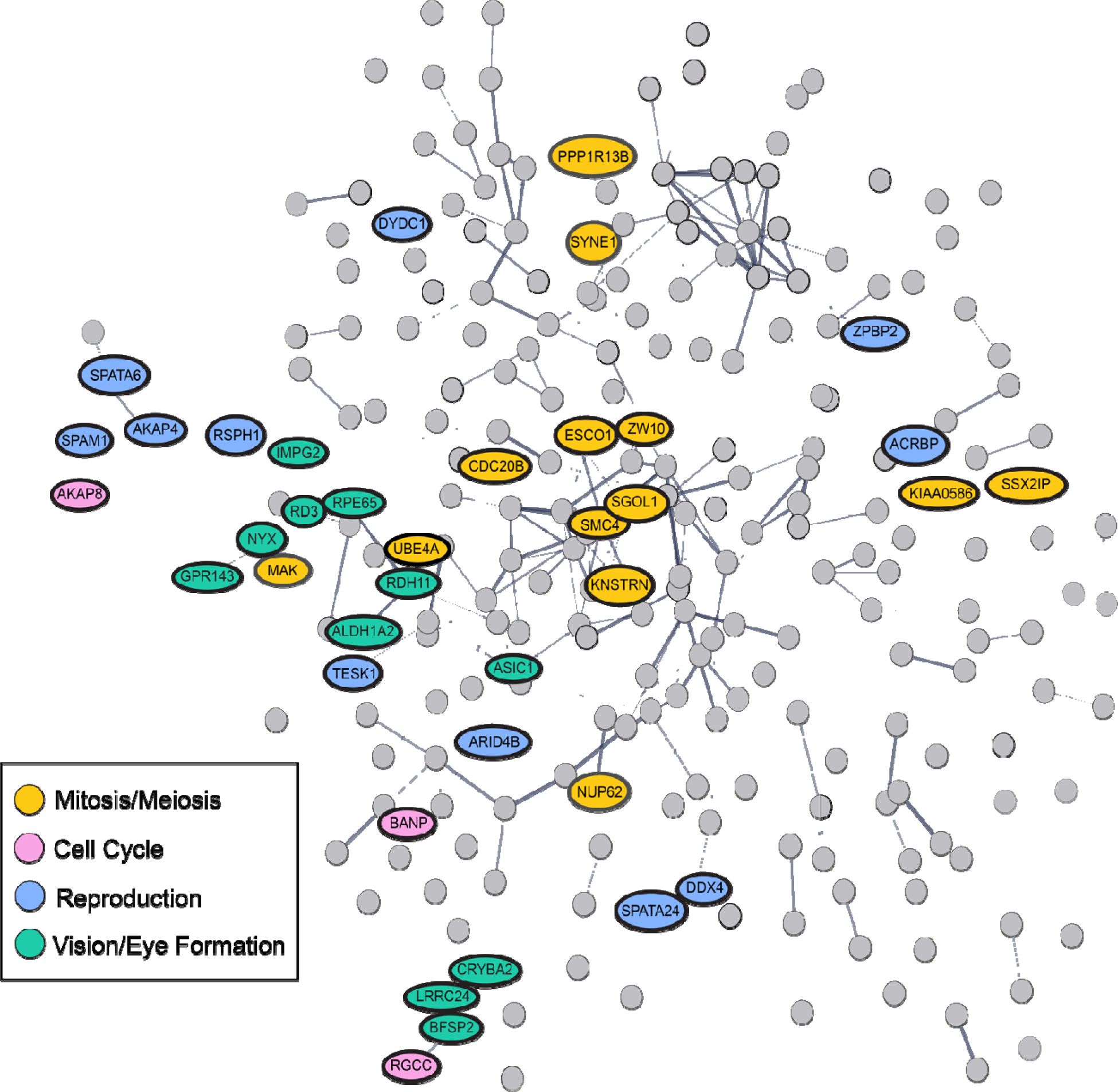
String network of genes under PS between *G. morafkai* and *G. agassizii*. The identified genes are grouped according to related functionality of chromosomal functions, cell cycle, reproduction, or vision and eye formation.

**Table 1:** Genome Assembly Statistics. Assembly statistics for the *G. morafkai* genome including length, scaffold length statistics of L50/N50 and L90/N50, number of total scaffolds and largest scaffold length in base pairs.

|  |  |
| --- | --- |
| Total Length (bp) | 2,221,617,999 |
| L50 | 47 |
| N50 (bp) | 14,713,492 |
| L90 | 173 |
| N90 (bp) | 2,550,679 |
| Number of Scaffolds | 1,557 |
| Largest Scaffold (bp) | 67,752,893 |

**Table 2:**
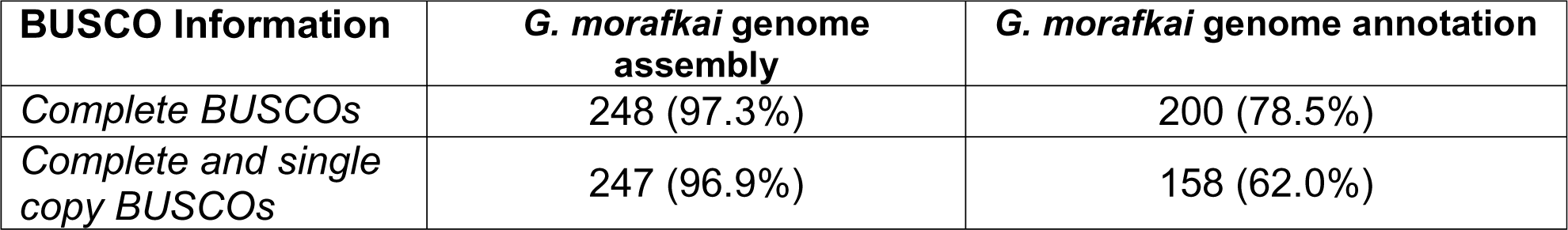
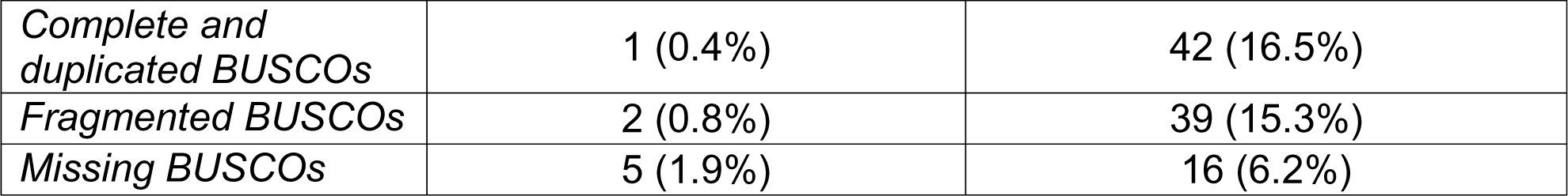
Genome Assembly BUSCO Statistics. BUSCO results from *G. morafkai* genome assembly and annotation, showing complete BUSCOs, complete and single copy BUSCOs, complete and duplicated BUSCOs, fragmented BUSCOs, and missing BUSCOs.

| BUSCO Information | *G. morafkai* genome assembly | *G. morafkai* genome annotation |
| --- | --- | --- |
| Complete BUSCOs | 248 (97.3%) | 200 (78.5%) |
| Complete and single copy BUSCOs | 247 (96.9%) | 158 (62.0%) |
| *Complete and duplicated BUSCOs* | 1 (0.4%) | 42 (16.5%) |
| *Fragmented BUSCOs* | 2 (0.8%) | 39 (15.3%) |
| *Missing BUSCOs* | 5 (1.9%) | 16 (6.2%) |

**Table 3:** Gene enrichment g:Profiler results. GO terms and IDs for enriched biological processes from genes under PS between *G. morafkai* and *G. agassizii*

| GO Terms | GO IDs | Adjusted p-value |
| --- | --- | --- |
| cellular component organization or biogenesis | GO:0071840 | 2.25E-08 |
| organelle organization | GO:0006996 | 1.28E-07 |
| cellular component organization | GO:0016043 | 2.33E-07 |
| organonitrogen compound metabolic process | GO:1901564 | 3.01E-07 |
| localization | GO:0051179 | 9.43E-05 |
| organelle assembly | GO:0070925 | 1.01E-04 |
| macromolecule localization | GO:0033036 | 8.83E-04 |
| regulation of cellular component organization | GO:0051128 | 1.48E-03 |
| establishment of localization | GO:0051234 | 2.32E-03 |
| regulation of catalytic activity | GO:0050790 | 2.41E-03 |
| cellular localization | GO:0051641 | 2.90E-03 |
| regulation of biological process | GO:0050789 | 3.79E-03 |
| protein metabolic process | GO:0019538 | 4.03E-03 |
| regulation of organelle organization | GO:0033043 | 4.05E-03 |
| biological regulation | GO:0065007 | 4.06E-03 |
| macromolecule modification | GO:0043412 | 5.52E-03 |
| establishment of protein localization | GO:0045184 | 6.51E-03 |
| protein localization | GO:0008104 | 6.63E-03 |
| cellular macromolecule localization | GO:0070727 | 7.31E-03 |
| cellular response to stress | GO:0033554 | 8.75E-03 |
| DNA damage response | GO:0006974 | 8.95E-03 |
| cytoskeleton organization | GO:0007010 | 1.48E-02 |
| regulation of molecular function | GO:0065009 | 1.52E-02 |
| transport | GO:0006810 | 2.06E-02 |
| regulation of metabolic process | GO:0019222 | 2.68E-02 |
| regulation of cellular process | GO:0050794 | 2.84E-02 |
| positive regulation of biological process | GO:0048518 | 3.58E-02 |
| protein modification process | GO:0036211 | 3.62E-02 |
| response to stimulus | GO:0050896 | 4.38E-02 |
| DNA metabolic process | GO:0006259 | 4.52 E-02 |
| regulation of macromolecule metabolic process | GO:0060255 | 4.55 E-02 |

### STRING Network Analysis of Positively Selected Genes

#### Genes Involved in Mitosis, Meiosis, and Chromosome Segregation

Chromosome-related processes central to mitosis and meiosis (GO:0051321, GO:0000278, GO:0007049, GO:0000819) were found to be under PS and many formed interacting subnetworks within the graph. Genes related to kinetochore development such as the Zw10 Kinetochore protein (*ZW10*), which functions in the proper segregation of chromosomes during cell division, and another kinetochore gene, *KNSTRN* (Kinetochore Localized Astrin Binding Protein) were both positively selected. Chromatid-related PS genes were *ESCO1*, which ensures proper sister chromatid cohesion, Shugoshin 1 (*SGOL1*), which protects centromeric cohesin proteins of sister chromatids to prevent mis-segregation, and *MAK*, which contributes to maintaining the synaptonemal complex that pairs homologous chromosomes during meiosis. Genes associated with the structure of chromosomes associated with chromosome segregation and condensation were *SMC4* and *UBE4A*.

#### Genes Involved in Cell Cycle Progression

Genes associated with cell cycle progression (GO:0007049) under PS were *RGCC, BANP*, *and AKAP8*. Interestingly, the *RGCC and BANP* genes also play a role in the p53 pathway, associated with tumor suppression and senescence, suggesting interaction of genes interfacing between cell cycle regulation associated with senescence and tumor suppression are under selection^24–26^. *AKAP8* assists with facilitating cell cycle progression and is needed for transition points in the cell cycle^27^.

#### Genes Involved in Reproduction

Reproductive genes under PS were associated with spermatogenesis evolution and sperm-egg binding processes (GO:0019953, GO:0007283, GO:0035036, GO:0007339). Interactions were identified between PS genes relating to central radial structures of the sperm flagellum (*RSPH1*), formation of the connecting piece between flagellum and sperm head (*SPATA6*), and structural component of the fibrous sheath affecting flagellum motility (*AKAP4*)^28^. Other genes that are localized to the testes are *ARID4B* and *DDX4*. *ARID4B* is expressed in the testes and serves a role in the blood-testis barrier in mice^29^. *DDX4* is typically expressed in spermatocytes and spermatids^30^. Genes involved in formation of the membraned organelle (the acrosome) at the tip of the sperm head that contains enzymes to allow penetration of the egg’s jelly-like coating (the zona pellucida) were under PS as well, specifically *DYDC1* and *ACRBP*, involved in forming the acrosome. *SPAM1* (Sperm Adhesion Molecule 1) and *ZPBP2* (Zona Pellucida Binding Protein 2) are involved both in acrosome formation and receptor signaling and adhesion of sperm to the zona pellucida of the egg. *TESK1* was another PS gene that plays a role in meiosis-related processes in spermatogenesis.

#### Genes Involved in Vision

There were nine PS genes involved in vision and eye formation that formed two interacting clusters (Fig 3). Cluster one had three genes involved in the ocular lens: Crystallin Beta A2 (*CRYBA2*), a main structural component of the eye lens, Beaded Filament Structural Protein 2 (*BFSP2*) which is a part of the lens and involved in optical clarity, and a Leucine Riche Repeat Containing 24 (*LRRC24*) synapse protein whose expression in humans is localized to bipolar cells, cone and rod photoreceptor cells^31,32^. A second cluster of six genes related specifically to photoceptors and retinal function. In this cluster were Retinoid Isomerohydrolase (*RPE65*) and Retinol Degeneration 3 (*RD3*)—both of which are critical to formation and/or maintenance of rod and cone photoreceptors, Interphotoreceptor Matrix Proteoglycan 2 (*IMPG2*) which organizes the matrix surrounding rod and cone photoreceptors, and Nyctalopin protein *(NYX*) that is thought to enhance signaling between bipolar and photoreceptive cells—both of which are implicated in vision under low-light conditions. Additionally, the G Protein-Coupled Receptor 143 (*GPR143*) localizes to melanocytes and plays a critical role in pigmentation and vision; defects in this gene cause ocular albinism^33^. Finally, Retinol Dehydrogenase 11 (*RDH11*) plays a role in activation of retinols and is involved in visual phototransduction. Together, these interacting PS genes indicate strong adaptation for photoreceptor function and light-dependent vision between *G. morafkai* and *G. agassizii*.

## DISCUSSION

The addition of the *G. morafkai* reference genome enables new analyses of the molecular mechanisms underpinning speciation of tortoises. Synteny analyses show broad conservation of genome-wide chromosomal structure between *G. morafkai* and *G agassizii*. A few chromosomal changes were identified between *G. morafkai* and *T. s. elegans,* supporting prior studies that synteny is highly conserved in chelonians^34,35^. This affirms chelonians may have not only low molecular evolution rates^36^ but low rates of chromosomal rearrangements, too.

The desert tortoises express subtle but clear and significant niche differentiation with some overlap potentially in the narrow hybrid zone where the species’ ranges overlap (black dots, Fig 2C). Most variation is explained by differences in precipitation patterns and temperature seasonality, showing *G. agassizii* residing in drier areas with more variable annual and seasonal precipitation, and *G. morafkai* resides in habitats with more precipitation and less temperature seasonal variation than *G. agassizii*, consistent with existing knowledge of the Mojave and Sonoran Deserts^37–39^. Similar niche specialization has been found in other xeric snake and lizard species. For example, in the neighboring regions of the Sonoran and Chihuahuan deserts, one study found evidence for population divergence in several snake species largely driven by climate differences between the two desert regions^40^. Another study focusing on *Crotalus* rattlesnakes found evidence that isolation by refugia in the Pleistocene era in the Mojave and Sonoran deserts may have contributed to desert adaptations of the rattlesnake species endemic to these areas^41^6/6/2024 11:50:00 AM. A species of Australian skink*, Liopholis kintorei*, also displays evidence for localized adaptation to temperature and precipitation differences across its range^42^. Such examples show that specialized niche use and environmental adaptation have been characterized in a number of reptile species adapted to desert environments. In this study we were able to document the genes underlying such ecological differences between two desert-adapted tortoises.

*Gopherus morafkai* and *G. agassizii* displayed extensive genomic reinforcement mechanisms, including chromatid pairing and segregation, sperm and egg formation and binding pathways, and cell cycle processes under PS. Another process under PS is vision development, which could be signaling the potential for an environmentally driven adaptive divergence response to the differential UV levels of the two deserts (Figure 2B). The UV differences could be tied to the evolution in lens development of the eyes of these species. Evolution in genes associated with low light conditions is also present and under PS, and could be a signal of the effects of contrasting behaviors between the tortoises during the winter brumation time, when *G. agassizii* individuals tend to stay in their dark, underground burrows, and *G. morafkai* is basking and readily exposed to sunlight^19^.

One of the strongest positive selection signals were in genes related to cell cycle and associated physical pairing and segregation of chromosomes relevant for mitosis and meiosis. One major mechanism under PS is the Spindle-Assembly Checkpoint (SAC), which is an active signaling point during the metaphase-anaphase transition of meiosis and mitosis. Chromosomes must be attached via microtubules to the spindles and kinetochores to ensure proper segregation of sister chromatids. Dynein motor proteins help align homologous chromosomes; once the requisite tension has accrued on the microtubules the SAC is triggered and transition to anaphase begins. The SAC has been proposed as a key mechanism of chromosomal reinforcement^43^. It is intuitive how structural chromosomal changes would cause SAC failure, but our findings indicate functional evolution in SAC-related kinetochore (*ZW10*), spindle (*SPDL1*), dynein (*BICD1, BICD2*), and chromosome-binding (*SGOL1*) genes imply SAC divergence may play a key role in genomic reinforcement even among highly syntenic lineages where large chromosomal structural changes are absent. The SAC and kinetochore related genes have been previously identified to be undergoing evolution in primates and other species^44,45^. Genes generally associated with mitosis, meiosis, the cell cycle, and chromosomal functions have been found repeatedly under selection broadly in other taxa such as *Drosophila*, *Arabidopsis*, and ant species; here we expand the empirical support for this form of postzygotic reinforcement to include reptiles as well^46–48^. Related observations include genes involved in crossing over in female *Drosophila*^46^ and proper chromosome condensation and cohesion in meiosis in *Arabidopsis*^49^.

Cell cycle progression was also under PS in *RGCC*, *BANP*, and *AKAP8*. These genes are also part of the p53 pathway, noted for its involvement in senescence and aging^50^. Tortoises are known for being long-lived reptiles with a large range of lifespans (∼20-60 years depending on species^22,51^, with most wild adults not living past 50 years^52^) and differing generation times (∼13-20 years)^53,54^. Literature suggests *G. morafkai* and *G. agassizii* have different lifespans, with *G. morafkai* living longer in the wild than *G. agassizii*^22^. This, combined with evidence that while most turtle species have long lifespans, they also have lower cancer rates^55^ and may lack senescence^56^, could potentially signal a unique balancing act of continuous growth and cell cycle control to avoid mutations, genomic deterioration, and senescence by selection acting on cell cycle processes.

Reproductive functions were under PS, as expected. Our results revealed many PS genes were related to sperm function and movement as well as sperm-egg recognition. Female tortoises can store sperm for use when ovulation occurs in the spring months. Previous studies have noted that long-term sperm storage occurred for upwards of two years in some *G. agassizii* females^57^. While there is little knowledge about use of sperm storage in mate choice and sperm competition in *Gopherus* species, other chelonian species use sperm storage competitively^58^. In this case, selection could be acting on the genes for sperm motility in the context of sperm competition^59^ between these species. This mechanism of sperm competition is common in other taxa such as birds^60^, squamates^61^, and even some species of bats^62^. Another process under PS was sperm-egg recognition, specifically formation of the sperm acrosome (*DYDC1*, *ACRBP,* and *SPAM1)*, and adhesion of sperm to the egg’s zona pellucida (*ZPBP2*). The sperm-egg interactions can directly underly species recognition, and we interpret this here as direct evidence of the evolution of prezygotic isolation mechanisms in these species^63^.

Genes under PS with ecological relevance related to eye and vision development, particularly photoreceptor and phototransduction under low light conditions. The adaptive qualities in these vision developmental processes could stem from behavioral differences in activity levels and circadian rhythm between the two species. *Gopherus morafkai* is known to occasionally leave their burrows in the winter periodically to bask in the sun and drink, while *G. agassizii* typically remains underground, in lower light conditions than *G. morafkai*, for the whole of the brumation period^19^. The differences in UV exposure across the two deserts could also prove to be a factor in selection acting on vision development. Previous studies have noted the possibility of lens evolution occurring as a stress response to avoid UV retinal damage^64^, as well as the functionality of the eye lens and cornea in absorbing UV light to avoid reaching the retina^65^. Other turtle species see different wavelengths of light, such as UV wavelengths^66^, which could suggest that PS genes associated with the development of rods and cones in the eyes could be an adaptation to the UV index transition between the deserts.

Additional themes such as mitochondrial-nuclear interactions, DNA replication, and immune system functionality were under PS between *G. morafkai* and *G. agassizii* (Supplemental Information). The mitochondrial-nuclear interactions could demonstrate possible discordance of the maternal mitochondrial genomes and nuclear genomes of hybrid tortoise individuals^67^, demonstrating another example of postzygotic reinforcement. PS in genes related to DNA replication and genome maintenance and stability could also reinforce lineage divergence through genome instability occurring in hybrids with incompatible DNA replication mechanisms^68,69^. Rapid evolution of the immune system is well-established as host defenses must evolve against new pathogens and there were several positively selected genes that have a variety of immune functions between *G. morafkai* and *G. agassizii*^70^. The immune genes under PS show evidence for both the innate and adaptive immune systems of these species undergoing differential adaptations to novel pathogens.

In conclusion, we present a chromosomal-scale *G. morafkai* reference genome and through comparative genomic analyses reveal an extensive array of genes that are predicted by theory to underlie core prezygotic and postzygotic reinforcement mechanisms of speciation, such as the spindle assembly checkpoint, chromatid pairing, and sperm-egg binding. Adaptation in vision processes under PS could be responding to environmental differences in UV exposure across the desert regions that *G. morafkai* and *G. agassizii* reside in, while other environmental differences were noted as significant in the environmental niche analysis. The analyses of synteny reveal few chromosomal changes between the closely related desert tortoise species and several genomic structural changes between *G. morafkai* and *T. s. elegans,* showing the genomic similarities between species within the *Gopherus* genus and differences with an ancestral turtle relative.

## ONLINE METHODS

### Genome Sequencing and Assembly

The type individual was an adult male collected in 2016 at 32° 28’ 39.5” N, 111° 07’ 21.2” W near Tucson, AZ. Skeletal muscle was obtained from the freshly euthanized carcass under Arizona State University’s Institutional Animal Care and Use Committee (IACUC) protocol #16-1508R.

#### PacBio Library and Sequencing

DNA samples were quantified using Qubit 2.0 Fluorometer. The PacBio SMRTbell library (∼20kb) for PacBio Sequel was constructed using SMRTbell Express Template Prep Kit 2.0 using the manufacturer recommended protocol. The library was bound to polymerase using the Sequel II Binding Kit 2.0 (PacBio) and loaded onto PacBio Sequel Sequencing was performed on PacBio Sequel II 8M SMRT cells generating 178 Gb of data.

#### Wtdbg2 Assembler

Wtdbg2^71^ was run to generate a primary assembly. Blobtools v1.1.1^72^ was used to identify potential contamination in the assembly based on blast (v2.9)^73^ results of the assembly against the NT database. A fraction of the scaffolds was identified as contaminant and were removed from the assembly. The filtered assembly (filtered.asm.cns.fa) was then used as an input to purge_dups v1.1.2^74^ and potential haplotypic duplications were removed from the assembly, resulting in the final purged.fa assembly.

#### Dovetail Hi-C library preparation and sequencing

A Dovetail Hi-C library was prepared in a similar manner as described previously^75^. Briefly, for each library, chromatin was fixed in place with formaldehyde in the nucleus and then extracted fixed chromatin was digested with DpnII, the 5’ overhangs filled in with biotinylated nucleotides, and then free blunt ends were ligated. After ligation, crosslinks were reversed and the DNA purified from protein. Purified DNA was treated to remove biotin that was not internal to ligated fragments. The DNA was then sheared to ∼350 bp mean fragment size and sequencing libraries were generated using NEBNext Ultra enzymes and Illumina-compatible adapters. Biotin-containing fragments were isolated using streptavidin beads before PCR enrichment of each library. The libraries were sequenced on an Illumina HiSeq X to produce 474 million 2x150 bp paired end reads.

#### Scaffolding the assembly with HiRise: Scaffolding with Chicago and HiC HiRise

The input de novo assembly, Chicago library reads, and Dovetail HiC library reads were used as input data for HiRise, a software pipeline designed specifically for using proximity ligation data to scaffold genome assemblies^76^. An iterative analysis was conducted. First, Chicago library sequences were aligned to the draft input assembly using a modified SNAP^77^ read mapper. The separations of Chicago read pairs mapped within draft scaffolds were analyzed by HiRise to produce a likelihood model for genomic distance between read pairs, and the model was used to identify and break putative misjoins, to score prospective joins, and make joins above a threshold. After aligning and scaffolding Chicago data, Dovetail HiC library sequences were aligned and scaffolded following the same method.

### Genome Annotation

For the annotation, we modeled species-specific repeats with RepeatModeler v2.0.1^78^ and generated a genome-guided transcriptome from mRNA data collected from three conspecific individuals in Edwards et al., 2016 (SRAs: SRS917892, SRS918084, SRS918109). For the transcriptome, we ran FASTQC v0.11.8^79^ for quality control and then corrected and filtered the reads using Rcorrector v1.0.4^80^ and TranscriptomeAssemblyTools^81^ and dynamically trimmed corrected reads using TrimGalore! V0.6.5^82^. Using Gmap and Gsnap v2020-06-30^83^ we mapped reads to the reference genome and converted the output into sorted bam files with Samtools v1.9^84^. To assemble the transcriptome we used Trinity v2.11.0^85^: --no_salmon genome_guided_maxintron 10000.

The species-specific repeat library and transcriptome assembly were inputted to Maker v3.01.03^86^ as repeat and EST evidence, respectively. In addition, for protein evidence we used reviewed proteins of human and chicken downloaded from UniProt (accessed August 10, 2020) and the predicted proteins for *Chrysemys picta bellii* from NCBI GCF_000241765.3. Round one of MAKER inferred gene models directly from these sources of evidence, followed by two rounds of *ab initio* gene prediction using SNAP v2006-07-28 and Augustus v3.2.3^87^. Gene models with an annotation edit distance (AED) of 0.25 or lower and a length of at least 50 were taken from round one and validated with fathom v2013-02-16 and any gene models with errors were removed and the clean file was used to generate the Hidden Markov Model in SNAP. To train Augustus, we split gene models into training and testing sets and followed standard series of training and optimizing of the gene models for this species. This training was performed before round two and three of MAKER.

Using accessory scripts from MAKER, we performed blastp with Swiss-PROT dataset (accessed 30 Dec 2020) (evalue 1e-6 -outfmt 6) and interproscan (appl pfam - dp -f TSV -goterms -iprlookup -pa -t p) to provide functional annotations to the gene models in the annotation file and predicted protein and transcript files. Finally, we pared down the final annotation file using the –clean function in GFFcleaner v0.10.1. We assessed gene completeness of the genome assembly, the genome-guided transcript, and the final annotation file using BUSCO v4.1.4^88^ and the eukaryote (eukaryote_odb10) dataset.

### Synteny Analysis

Synteny was analyzed across the species using SynChro^89^ and CHROnicle^89^.To extract proteins from the genome assembly and annotation files, a modified version of gff2fasta.pl script from ISUgenomics^90^ was used. The top 30 gene-rich scaffolds with the largest number of genes were subset from each species using scripts from Biopython v1.73^91^. The protein files were then formatted for SynChro input using the CHROnicle ConvertFasta.py script. The SynChro.py script was run pairwise on the formatted files to produce the syntenic blocks and svg images. A delta value of 2 was used in this step, specifying the number of orthologous genes to be syntenic between species to be formed as a syntenic block. The chromosomal painting files were then ordered by descending scaffold size as sorted in the genome assembly. Each scaffold was scaled down to a smaller size using the base pair size from the genomic assembly to view the correct size of each scaffold.

### dN/dS Analysis

dN/dS was calculated with the dNdS argument from the orthologr v0.4.0^92^ package. Orthologs from CDS sequences generated using GffRead v 0.12.7 from each species were used in a pairwise comparison and dN/dS ratios were computed using the well-known Comeron’s and Li’s methods. The default method orthologr uses is Comeron’s method, which is a combination of several methods of calculation. Genes with a dN/dS ratio of 1.0 and above were identified as under PS from the full output of orthologr, and were used for downstream genomic analyses. The genes under PS for the pairwise comparison from each method of calculating dN/dS were then intersected to generate the final set of genes identified as under PS from both methods of calculation selection.

### Gene Set Enrichment Analysis

The genes identified as under PS in each pairwise comparison were input in g:Profiler e109_eg56_p17_1d3191d^94^ to identify any potential significant enrichment of biological processes among the gene set. The enrichment analysis was run using each list of positively selected genes against the *Mus musculus* genome due to its high content of known annotations from Ensembl databases. The threshold used in the enrichment analyses was the default g:SCS threshold setting from g:Profiler with a significance threshold of 0.05. The output enrichment files for biological processes were used as input for REVIGO v1.8.1^95^ in order to produce treemap images of the biological processes identified as enriched. The processes were grouped into a “supercluster” by color and the size of each cluster is representative of p value. Smaller clusters in the treemap were further grouped into a single cluster named by the largest representative GO category in the original cluster. See Supplemental Figure 6 for the full treemap and detailed superclusters.

### Gene Interaction Analysis

To determine whether the genes under PS interacted with one another, we took the final set of positively selected genes and made a network of known interactions among them using the String Database v11.5^96^. Analyses used human as a reference organism; edges were weighted by confidence of interaction and no nodes were added to the network that was not on the list of positively selected genes. In order to group genes by functionality into clusters, GeneCards and Gene Ontology’s AmiGO database AmiGO 2 2.5.17^97^ were utilized for each gene’s definition, as well as related GO categories that could define over larger groups.

### Tortoise Distribution Map

Tortoise occurrence data was collected from the Global Biodiversity Information Facility (GBIF)^98^ and used as input in QGIS v3.22.1^99^. Polygons were created based on this occurrence data to form the tortoise distributions range map.

### Tortoise Climatic Niche Characterization

Georeferenced records were obtained for the *Gopherus* species from the GBIF database. Occurrences were manually curated and deduplicated within 1 km to accurately represent each species distribution. Then, climatic data was obtained from Worldclim^100^ for 19 bioclimatic variables derived from temperature and precipitation (Table S1), with 1 km of spatial resolution. From each occurrence record the values for all bioclimatic variables were extracted. To characterize the climatic niche of the four species, a principal component analysis (PCA) was performed with the bioclimatic data in R 4.1.2^101^. Niche divergence was assessed by estimating niche overlap with the “ellipsenm” package^102^ in R, using the 19 bioclimatic variables. Significance was obtained from 1,000 bootstrap replicates.

### UV Index and Map

Monthly UV index data with a 0.5 degree resolution from 2010 was obtained from NASA Earth Observations^103^. Then, the UV index annual average was calculated and applied to a map using QGIS.

## ACKNOWLEDGEMENTS

The authors acknowledge Research Computing at ASU for providing high performance computing resources that have contributed to the research results reported within this paper.

## DATA AVAILABILITY

The supporting genome and annotation files are deposited in the Harvard dataverse. The genome assembly is available under NCBI bioproject <u>PRJNA825533</u>.

